# Hypo-osmotic Stress Induces ATP Release via Volume-regulated Anion Channels in Undifferentiated Mammary Cells

**DOI:** 10.1101/2021.04.25.441329

**Authors:** Kishio Furuya, Yuko Takahashi, Hiroaki Hirata, Takeshi Kobayashi, Mikhail Samsonov, Masahiro Sokabe

## Abstract

The high interstitial ATP concentration in the cancer microenvironment is a major source of adenosine, which acts as a strong immune suppressor. However, the source of ATP release has not been elucidated. We measured the ATP release during hypotonic stress using a real-time ATP luminescence imaging system in primary cultured mammary cells and in breast cell lines. In primary cultured cells, ATP was intermittently released with transient-sharp peaks, while in breast cell lines ATP was released with a slowly rising diffuse pattern. The diffuse ATP release pattern was changed to a transient-sharp pattern by cholera toxin treatment and the reverse change was induced by transforming growth factor (TGF) β treatment. DCPIB, an inhibitor of volume-regulated anion channels (VRACs), only suppressed the diffuse pattern. The inflammatory mediator sphingosine-1-phosphate (S1P) induced a diffuse ATP release pattern isovolumetrically. The knockdown of A isoform of leucine-rich repeat-containing protein 8 (LRRC8A), the essential molecular entity of VRACs, using shRNA suppressed the diffuse pattern. These results suggest that abundantly expressed VRACs are a conduit of ATP release in undifferentiated cells, including cancer cells.

## Introduction

Extracellular ATP is a ubiquitous mediator of local intercellular signaling within the body (Burnstock and Virgilio, 2013). Extracellular ATP is quickly hydrolyzed by ecto-ATPases and is maintained at a concentration near zero in the interstitial fluids of unstressed tissues. However, the ATP concentration is considerably high at sites of inflammation or in cancerous tissues (Pellegatti P et al., 2008; Virgilio FDi, 2012) despite the abundance of ecto-ATPase. The functions of extracellular ATP are gradually being clarified. One major characteristic of cancer is the suppression of the immune attack on tumor cells in the cancer microenvironment. This makes cancer immune therapy difficult. A chronically increased level of adenosine was shown to contribute to this immunosuppression via multiple pathways, including the inhibition of T cells and dendritic cells (Antonioli L et al., 2013a). A major source of adenosine is the hydrolysis of ATP by the ecto-ATPases, CD39 and CD73. CD39 is expressed in regulatory T cells (Antonioli L et al., 2013b), and both enzymes exist abundantly in the cancer microenvironment. These enzymes are now considered to be therapeutic targets (Hausler SF et al. 2014). The adenosine pathway to suppress antitumor immune responses also affected the efficiency of immunotherapy in a recent clinical trial using anti-PD-1/PD-L1 monoclonal antibody (mAbs) as immune checkpoint inhibitors (Beavis PA et al., 2015b). Furthermore, adenosine receptor 2A blockade on T cells significantly enhanced the efficacy of anti-PD-1 mAb and increased the survival of mice inoculated with CD73^+^ tumors (Beavis PA et al., 2015a).

Despite increasing evidence to support the importance of adenosine pathways in the cancer microenvironment, the source of ATP, which is itself a major source of adenosine, remains unclear. The existence of high concentrations of ATP at sites of inflammation or in the cancer microenvironment is readily accepted due to the presence of destroyed or dying cells. However, these cells do not release much ATP and/or the release is not sustained. To maintain a high ATP concentration in the cancer microenvironment, ATP must be continuously released in a regulated manner; however, the mechanisms through which this occurs are not known. In the present study, we revealed the co-existence of two different patterns of ATP release following hypotonic stress in mammary epithelial cells using our real-time ATP luminescence imaging microscope. One was a diffuse prolonged pattern of ATP release, that was usually observed in breast cell lines but not in primary cultured normal mammary epithelial cells. This diffuse ATP release pattern was blocked solely by DCPIB, a blocker of volume-regulated anion channels (VRACs) (Decher N et al., 2001; Friard J et al., 2017), suggesting the involvement of these channels in the diffuse release of ATP.

VRACs are a major mechanism of vertebrate cell volume regulation and also participate in numerous physiological and pathophysiological processes, including cancer, edema, cell proliferation, migration, angiogenesis and apoptosis (Okada Y et al., 2009; Pedersen SF et al., 2016). The cell volume dramatically changes with the cell cycle, and cell cycle progression is a cell-size dependent process (Ginzberg MB et al., 2015). VRACs are differentially regulated throughout the cell cycle, and the inhibition of VRACs suppresses proliferation in various types of cells, including hepatocytes, endothelial cells, smooth muscle cells and cancer cells (Pedersen SF et al., 2016). In 2014, leucine-rich repeat containing 8 family A (LRRC8A) was identified as an indispensable component of VRACs (Voss FK et al., 2014; Qiu Z et al., 2014), although at least one other family member is needed to mediate the VRAC current. LRRC8 is a distantly pannexin-1-related protein family and forms a hetero hexamer (Jentsch TJ et al., 2016; Abascal F and Zardoya R, 2012), which may be the reason for the extremely varied properties of VRACs. We herein confirmed the contribution of VRACs to the release of ATP by the knockdown of LRRC8A using shRNA.

In the present study we used hypotonic stress to stimulate the release of ATP. In higher organisms, systemic osmolality is kept constant by multilevel homeostatic control. Nevertheless, dramatic changes in osmolality occur in the kidney and gastrointestinal tract with the induction of the influx of large amount of saccharides, amino acids and sodium ions. Furthermore, cells experience frequent fluctuations in their volume due to unbalanced transmembrane fluxes of ions and nutrients or the synthesis and degradation of macromolecules within the cells. Under pathological conditions in the brain, ischemia, hyponatremia and epilepsy cause astrocytic edema (cell swelling) (Mongin AA, 2016). In cancer patients, the degradation of protein synthesis including albumin causes cachexia and edema (Fazzari J and Singh G, 2016). VRACs are also activated by various chemical stimuli, including sphingosine-1-phosphate (S1P). S1P is a signaling lysophospholipid and an inflammatory mediator like bacterial lipopolysaccharides, which abundantly exists in the cancer microenvironment. We herein report, for the first time, that prolonged diffuse ATP release via VRACs was induced by hypotonic stress and S1P application in undifferentiated breast cell lines, and that it was enhanced by treatment with transforming growth factor (TGF) β, a carcinogenic agent and reduced by treatment with cholera toxin, an anti-carcinogenic agent.

## Materials and Methods

### Cell culture

Mammary glands were dissected from lactating ICR mice (Japan SLC, Hamamatsu, Japan) after lethal deep anesthesia (pentobarbital Na or a combination anesthetic with medetomidine, midazolam and butorphanol). Mammary epithelial cells were isolated and cultured, as described previously (Nakano H et al, 1997) using Dispase II (Godo Shusei Co., Tokyo, Japan) and collagenase (Type III; Worthington, Freehold, NJ, USA).

#### Cell lines

The cancerous breast cell lines MCF7 (AKR-211, MCF-7/GFP, Cell Biolabs, Inc., San Diego, CA, USA) and MDA-MB231 (AKR-201, MDA-MB231/GFP, Cell Biolabs, Inc.) were cultivated in DMEM/F12 (Wako Pure Chemical, Osaka, Japan) supplemented with 10% FBS; the non-carcinogenic breast epithelial cell line MCF10A (CRL-10317, ATCC, Manassas, VA, USA) was cultivated on a collagen-coated dish in HuMEC (Gibco, Thermo Fisher Scientific, Waltham, MA, USA); at 37°C under 5% CO_2_.

For the measurements, cells of the primary culture and several cell lines were cultured on collagen gel (Cellmatrix type I-A, Nitta Gelatin, Osaka, Japan) on a 14- or 22-mmφ cover glass (#1, Matsunami Glass Ind. Ltd. Osaka, Japan) for 1 to 4 days at 37°C under 5% CO_2_, in sub-confluent to confluent conditions. In some experiments, 3 types of cells (e.g., different cell lines or different shRNA treated cells) each on 3 separated collagen-gel patches were cultured simultaneously on a 22-mmφ cover glass, which made it easy to compare the responses under the same conditions of cultivation and measurement. Cholera toxin (100 ng/ml) (Wako), cholera toxin B subunit (Wako) or TGF β (10 ng/ml) (Wako) was added to the culture medium for 3 h to 3 days before measurement in some cases.

### Experimental setup

The cells on the cover glass were set in a small perfusion chamber (approximately 100 μl in volume) on the stage of an upright microscope (BX51WI; Olympus, Tokyo, Japan) with a 1× (Plan UW, NA0.04; Nikon, Tokyo, Japan), 4× (340 Fluor XL, NA0.28; Olympus) or 10× (Plan Apo, NA0.45; Nikon) objective lens. The medium was replaced with DME/F12 buffered with 10 mM HEPES (pH 7.4) (Gibco) containing 40-50% luciferin-luciferase solution (see below). Medium changes (300 μl) were performed via capillary action for approximately 30 s without any mechanical effect of flow. Hypotonic solutions were made by adding a solution with 1.05mM CaCl_2_ + 0.7mM MgCl_2_ (30%-50% [v/v]), which keeps the Ca^2+^ and Mg^2+^ concentration constant in hypotonic solutions. The osmolality of each medium was DME/F12-HEPES: 303 mosm; 30% hypotonic solution: 218 mosm; 50%hypotonic solution: 155 mosm. In some cases, just distilled water was used to make hypotonic solutions, but no marked difference on the ATP response was observed. DCPIB (4-[(2-Butyl-6,7-dichloro-2 - cyclopentyl-2,3-dihydro-1 -oxo-1H-inden-5-yl) oxy] butanoic acid, TOCRIS, Bristol, UK), sphingosine-1-phosphate (Huzzah S1P, Human Serum Albumin/sphingosine-1-phosphate Complex; Avanti Polar Lipids, Inc., Alabaster, AL, USA) and other active reagents were added to the perfusion medium.

### Real-time imaging of the ATP release

The ATP release was measured in real-time using a luminescence imaging system that has been previously described (Furuya K et al., 2014). Luciferin-luciferase ATP bioluminescence was detected using a high-sensitivity camera system simultaneously with infrared-DIC imaging to monitor the cells. An osmolality adjusted luciferin-luciferase solution (Lucifer HS Set; Kikkoman Biochemifa Co., Tokyo, Japan; or Luciferase FM plus; Bioenex Inc., Hiroshima, Japan) was added to the perfusate with 40-50% volume. After standing for 15 min after a medium change, ATP-dependent luminescence was detected with a high-sensitivity EMCCD camera (Cascade 512F; Photometrics, Tucson, AZ, USA) equipped with a cooled image intensifier (C8600-04; Hamamatsu Photonics, Hamamatsu, Japan). Images were acquired at a frequency of 2 Hz with an exposure time of 500 ms using the MetaMorph software program (ver. 7.8; Molecular Devices, San Jose, CA, USA) in stream acquisition mode. For the data analyses, image smoothing was usually conducted by calculating the average of six sequential images. In our system, DCPIB (100 μM) itself does not affect luciferin-luciferase bioluminescence by ATP. ATP imaging experiments were performed at 30±2°C.

### Measurements of regulatory volume decrease (RVD)

It is known that VRACs are channels responsible for the regulatory volume decrease (RVD) that occurs after a volume increase by hypotonic stress. Volume changes during hypotonic stress (30%) were monitored by perpendicular cross-section image of the cells, which constitutively expressed GFP, using a confocal X-Z-T scan (LSM510, Carl Zeiss, Oberkochen, Germany) with a 63× NA1.4 objective, 4s scan speed and 10s interval. Volume change was estimated from a change in the fluorescence intensity of a certain unit area in the cross-section image and normalized by an image just before stimulation.

### Real-time polymerase chain reaction (qPCR)

The expression of LRRC8 family members (A to E) was measured by reverse transcription (RT) qPCR. mRNA specimens were isolated from the cells grown on collagen-gel in 24-well dishes using a NucleoSpin RNAplus RNA isolation kit (Macherey-Nagel, Dueren, Germany) and converted to cDNA with SuperScript IV VILO Master Mix (Invitrogen, Thermo Fisher Scientific, Waltham, MA, USA) or SuperScript III First-Strand Synthesis System (Invitrogen, Thermo Fisher Scientific). The expression was determined by RT-qPCR using a LightCycler 480 or Nano (Roche, Mannheim, Germany) with SYBR Green I Master Mix (Roche) and quantitative primers (Perfect Real Time Primer, Takara, Shiga, Japan, Supplemental Table S1). The expression was normalized to GAPDH within each sample.

### LRRC8A knockdown with shRNA

LRRC8A silencing with shRNA was performed using retrovirus mediated gene transfer as described previously (Hirata et al., 2017). To generate retrovirus expressing shRNA against LRRC8A, the target sequences (Supplemental Table S1) were inserted into the pSUPER.retro.puro retroviral vector (OligoEngine, Seattle, WA, USA). For control, a non-targeting sequence 5’-ATAGTCACAGACATTAGGT-3’ was introduced. The shRNA-containing vector was co-transfected with the pE-ampho vector into HEK293T cells using GeneJuice transfection reagent (Merck Millipore, Burlington, MA, USA). Supernatants containing viral particles were collected 48 h after the transfection, filtered through 0.45-μ m syringe filters, and used for infection into three breast cell lines in the presence of 8 μg/ml Polybrene (Sigma-Aldrich, St. Louis, MO). Infected cells were selected with 1.5 μg/ml puromycin (Sigma-Aldrich).

## Results

### Primary cultured cells and established cell lines showed different ATP release kinetics and patterns

Hypotonic (hypo-osmotic) stress was applied to induce the release of ATP from several types of mammary epithelial cells, and distinct differences between the cells in the primary culture and breast cell lines were revealed for the first time using our ATP-imaging system. In primary cultures of mammary epithelial cells from lactating mice, 30% hypotonic stimulation (70% osmolality, 218 mosm) induced the transient release of ATP, which occurred intermittently in several randomly distributed cells in the colony (Fig. 1A, Supplemental Movie 1; hereafter referred to as the “transient-sharp” pattern). The duration of each peak was several dozen seconds. In contrast, in a breast cell line (MDA-MB231), 30% hypotonic stimulation induced a slow-rising ATP release response with a diffuse appearance (Fig. 1B, Supplemental Movie 2; hereafter referred to as the “diffuse” pattern). In this pattern of release, we could not identify individual ATP-releasing cells. Interestingly, in primary cultured mammary cells, the local peak concentration of released ATP during each event was significantly higher than in the cell lines; however, the average ATP concentration in the entire area was greater and longer in the breast cell lines (Fig. 1Ac and Fig. 1Bc).

**Fig. 1.**
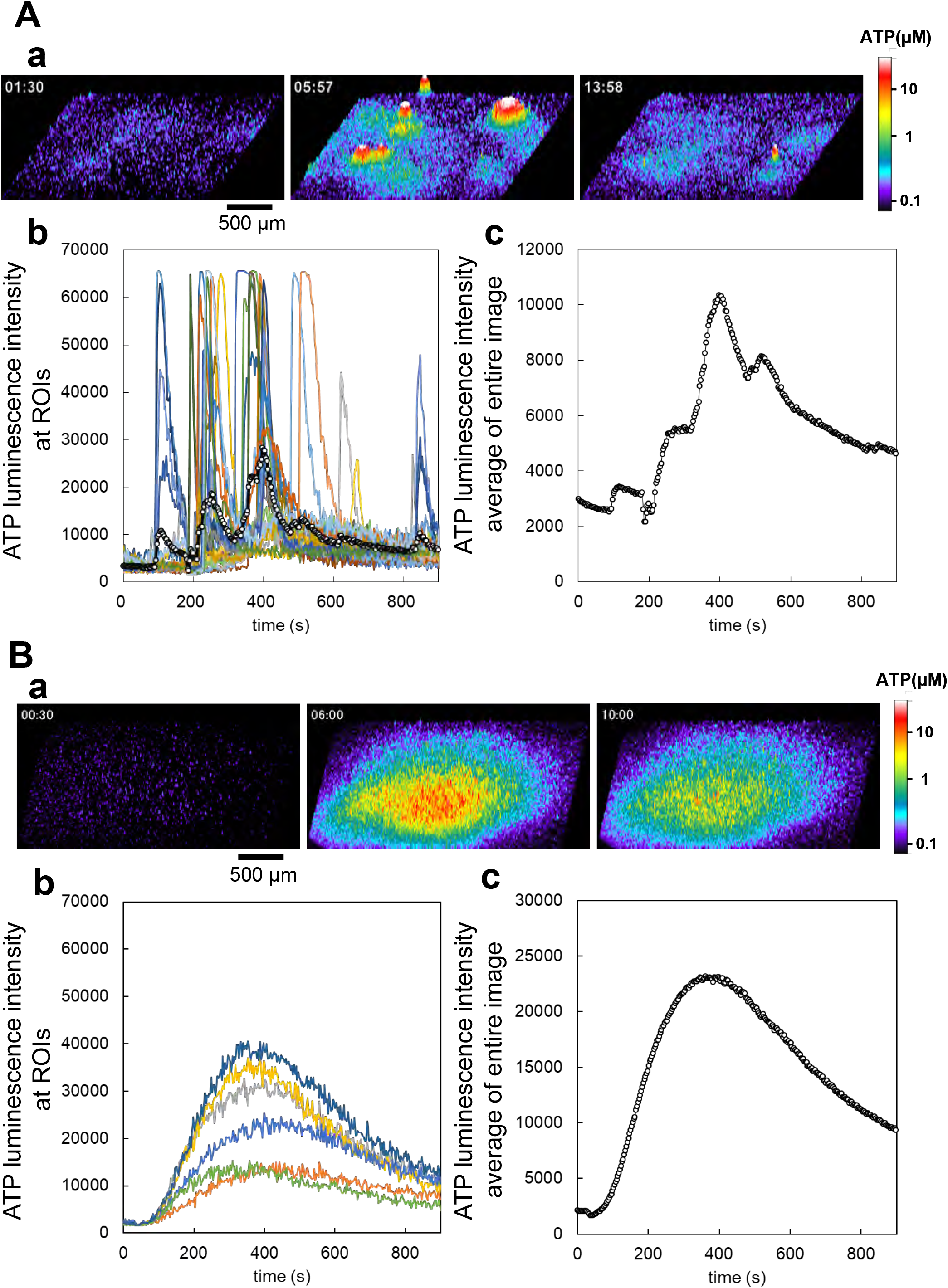
Real-time ATP luminescence imaging reveals different kinetics and patterns of ATP release induced by hypotonic stress in primary cultured mammary epithelial cells and breast cell lines. **A: a**, Hypotonic stress (30%, 70% osmolality, 218 mosm) induced transient ATP release events in primary mammary epithelial cells from a lactating mouse. Sharp responses occurred intermittently at different time points in several randomly distributed cells (“transient-sharp” response). The ATP release is shown in 3D-intensity profile (see also Supplemental Movie 1). **b**, The time course of luminescence changes in each responding cell. **c**, The time course of the average luminescence intensity in the whole observed area. The frequency of release events peaked at 200–400 s. **B: a**, Hypotonic stress (30%) induced diffuse ATP release in a breast cell line (MDA-MB231). The responses were slow-rising and had a diffuse appearance, such that ATP-releasing cells could not be identified (“diffuse” response). The ATP release is shown in 3D-intensity profile (see also Supplemental Movie 2). **b**, The time course of luminescence change observed at several points of cell culture. **c**, The time course of the average luminescence intensity in the whole observed area. The release of ATP peaked at 500– 700 s. Other breast cell lines (MCE7, MCF10A) showed similar diffuse responses following hypotonic stress.

Other breast cells lines, MCF7 carcinoma and MCF10A non-carcinogenic, and a human lung carcinoma cell line (A549) (Furuya K et al., 2014) exhibited a diffuse ATP release pattern. Conversely, other primary cultured cells from subepithelial fibroblasts in the rat small intestine (Furuya and Furuya, 2007) exhibited a transient-sharp ATP release pattern (Fig. S1).

### Cholera toxin alters the patterns of ATP responses from diffuse to transient-sharp

The two types of ATP responses were interchangeable. Diffuse ATP responses observed in MCF10A breast cell line (Fig. 2A, Supplemental Movie 3) or other carcinogenic cell lines (MCF7 and MDA-MB231) were modified to the transient-sharp pattern (Fig. 2B, Supplemental Movie 4) following cholera toxin treatment (100 ng/ml) for 7 h (range: 3 h to 3 days).

**Fig. 2.**
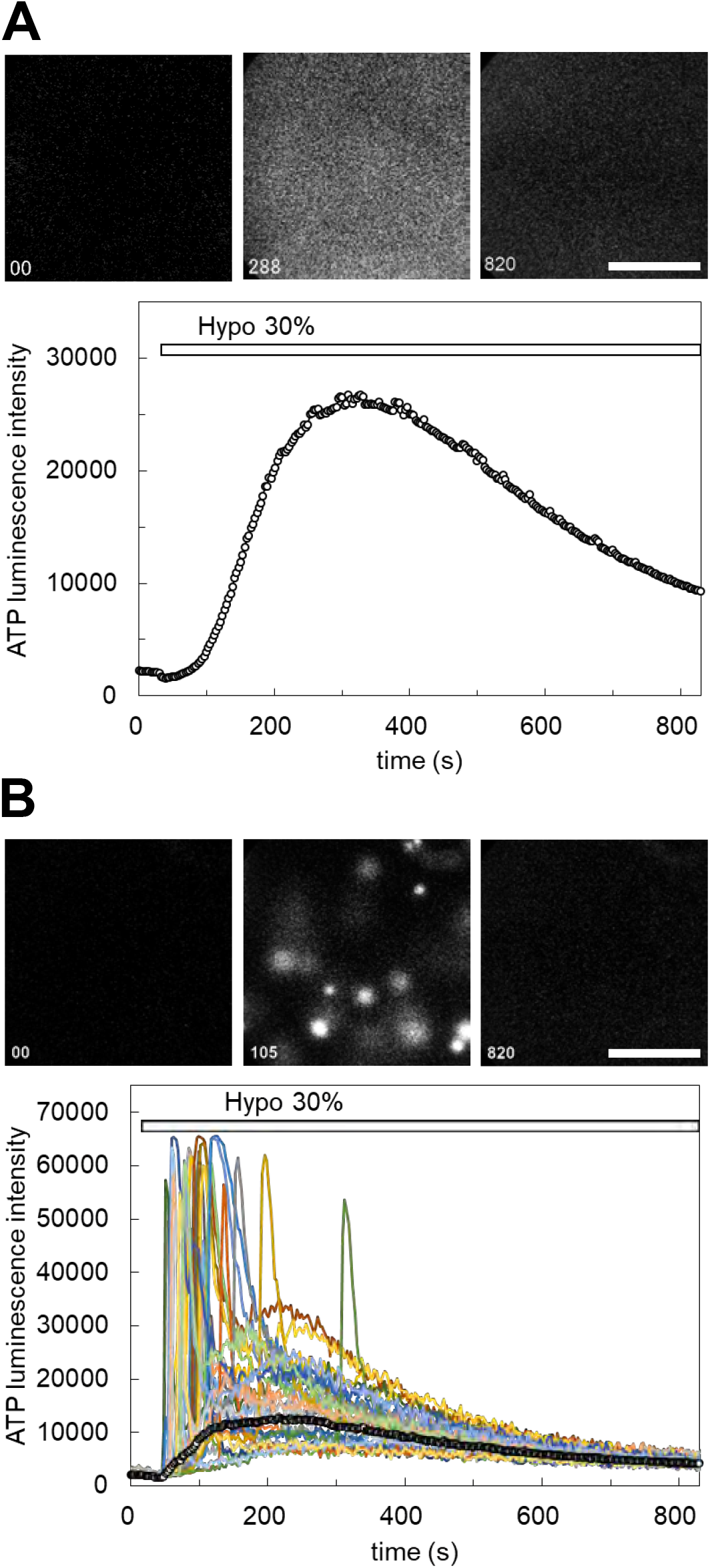
The pattern of ATP release was changed from “diffuse” to “transient-sharp” by a treatment with cholera toxin. **A:** Hypotonic stress (30%) induced slow-rising and diffuse ATP release in MCF10A cells. The release of ATP continued for a relatively long duration, and the decay of the ATP response was slow (see also Supplemental Movie 3). **B:** Cholera toxin (100 ng/ml) treatment for several hours changed the pattern of ATP release to the transient-sharp response type. Hypotonic stress (30%) was applied to MCF10A cells after 7-h cholera toxin treatment (see also Supplemental Movie 4). The other cell lines (MCF7, MDA-MB231) showed similar results. Scale bar indicates 500 μm.

The two types of ATP responses were not exclusive. Even in cholera toxin-untreated cells, transient-sharp ATP releases occasionally occurred in a small number of cells immediately after hypotonic stress overlapping with the diffuse pattern. In addition, spontaneous release of ATP with transient-sharp pattern was sometimes observed without hypotonic stress. The appearance of the transient-sharp pattern seemed to depend on the conditions of each cell in cell cultures; however the mechanisms through which this is induced remained to be elucidated.

### DCPIB blocks the diffuse release of ATP, but not the transient-sharp release of ATP

We assessed the following inhibitors of ATP release: CBX and 10PANX for hemi channels, NPPB and DCPIB for Cl^−^ channels, CFTR(inh)-172 for CFTR, and a cocktail of brefeldin A, monensin and NEM, and clodronate for exocytosis. Among them, only DCPIB, a specific inhibitor of VRACs (Decher N et al., 2001; Friard J et al., 2017), effectively inhibited the diffuse pattern. As shown in Fig. 3A and 3B, DCPIB treatment (100 μM) completely suppressed the diffuse ATP release pattern. The effects of DCPIB could be washed out by several medium changes over 15 min (Fig. 3B). However, DCPIB did not block the transient-sharp release pattern in cholera toxin-treated cells (Fig. 3C, Supplemental Movie 5) or in primary cultured cells. On the contrary, DCPIB treatment sometimes enhanced or induced a transient-sharp release of ATP.

**Fig. 3.**
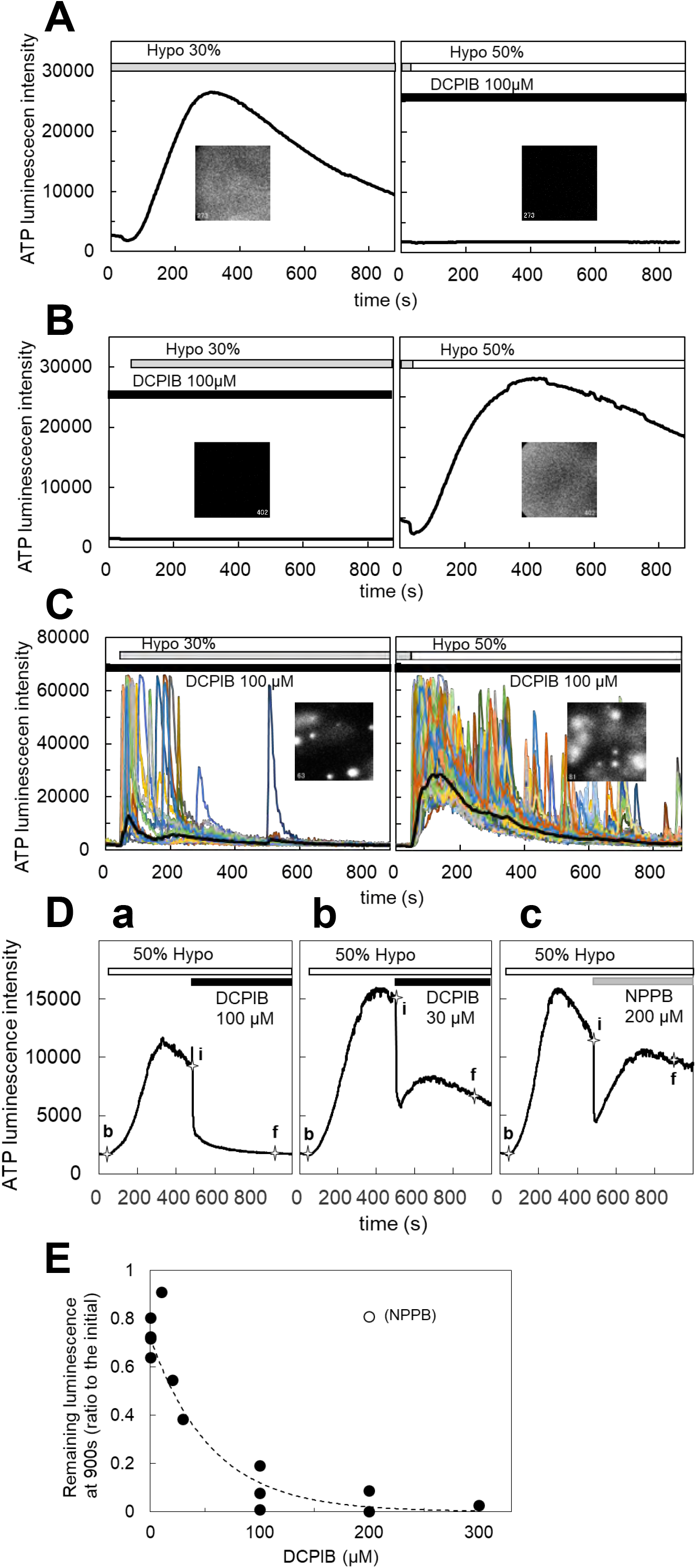
DCPIB, a blocker of the volume regulatory anion channels (VRACs), inhibited the slow diffuse response but not the sharp intermittent release. **A:** Hypotonic stress (30%) induced the diffuse ATP release pattern in MCF10A cells. DCPIB treatment (100 μM) suppressed the response to subsequent 50% hypotonic stress. **B:** DCPIB treatment (100 μM) suppressed the response to 30% hypotonic stress in MCF10A cells. After washout of DCPIB, the ATP response recovered with subsequent 50% hypotonic stress. **C:** The transient-sharp response in cholera toxin (100 ng/ml)-treated MCF10A cells was not affected by DCPIB treatment (100 μM) (see also Supplemental Movie 5). Other breast cell lines (MCF7, MDA-MB231) showed similar results. In these experiments (**A, B, C**), 50% hypotonic solution was applied sequentially over 30min after the application of a 30% hypotonic solution. **D:** The dose dependence of DCPIB in blocking diffuse ATP release were measured according to the level of decay of the diffuse ATP response after treatment with each concentration of DCPIB in MCF10A cells. The diffuse ATP release induced by 50% hypotonic stimulation (at 30 s; baseline intensity: b) was blocked by the perfusion of each concentration of DCPIB (at 480 s; initial intensity: i). The levels of decay were measured at 900 s (final intensity: f). **a**: DCPIB 100 μM. **b**: DCPIB 30 μM. **c**: NPPB 200 μM. The blocking effect of DCPIB at each concentration was calculated based on the value of (f-b)/(i-b). **E:** The dose dependence was determined based on the calculated value (f-b)/(i-b) at each concentration of DCPIB. The IC_50_ obtained by an exponential fitting curve was 38.5 μM. NPPB showed almost no effect, even at 200 μM.

The dose-dependent effects of DCPIB on the blockade of ATP release were measured according to the level of decay of the diffuse ATP response after treatment with each concentration of DCPIB (Fig. 3D). DCPIB blocked the release of ATP in a dose-dependent manner, while NPPB (200 μM) did not affect the release of ATP (Fig. 3D, a-c). The dose response curve (Fig. 3E) shows that the IC_50_ was 38.5 μM.

### S1P induced the diffuse release of ATP which was blocked by DCPIB

DCPIB blockade of the diffuse ATP response suggested that VRACs contributed to the diffuse release of ATP. VRACs are activated not only by hypotonic stress but also isovolumetrically by various intracellular factors. The inflammatory mediator sphingosine-1-phosphate (S1P) was reported to activate VRACs (Burow P et al., 2015). The application of S1P (100 nM-1 μM) induced the release of ATP with a slowly rising and diffuse pattern in MCF7 cells, which had a comparable amplitude to but declined somewhat faster in comparison to the response to 30% hypotonic stimulation (Fig. 4A). The responses to S1P (200 nM) and hypotonic stress (30%) were not competitive, rather they were additive (Fig. 4B, C). The response to S1P after hypotonic stimulation was enhanced in comparison to the sole administration of S1P (Fig. 4D). The ATP response to S1P was suppressed by DCPIB similarly to the response to hypotonic stress (Fig. 4E), suggesting that VRAC activation by S1P is involved in the diffuse ATP response.

**Fig. 4.**
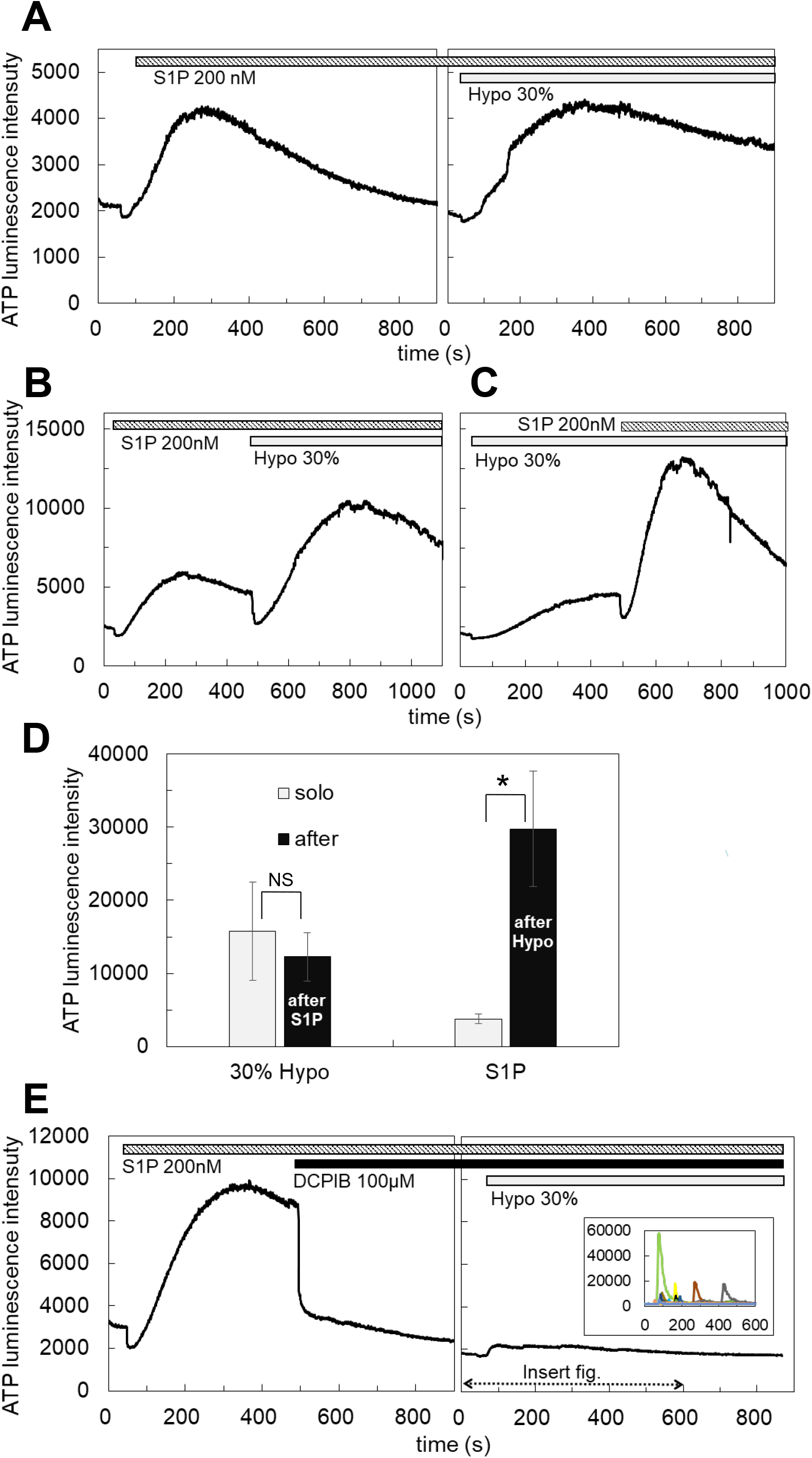
S1P induced the diffuse ATP release, which was blocked by DCPIB. **A:** The application of S1P (200 nM) induced the diffuse ATP release pattern in MCF7 cells, whose amplitude was comparable with the response to 30% hypotonic stimulation. **B, C:** The response to S1P was additive to the effect of 30% hypotonic stimulation. **D:** Especially the response to S1P after 30% hypotonic stress was significantly enhanced in comparison to the sole administration of S1P. Mean ± S.E. (N=6), t-test; NS: no significance, *: p<0.01. **E:** The response to S1P was suppressed by DCPIB similarly to the response to hypotonic stress. The small response induced by 30% hypotonic stimulation after DCPIB treatment was due to the average of several intermittent sharp responses, as shown in the insert figure, which were usually induced by hypotonic stress in DCPIB-treated cells.

### TGFβ enhanced the diffuse ATP release pattern induced by both hypotonic stress and S1P

Transforming growth factor (TGF) is critically important for mammary development and carcinogenesis, as it regulates various cellular activities, including the epithelial-mesenchymal transition (EMT) (Moses H and Barcellos-Hoff MH, 2011). In primary cultured mammary epithelial cells, hypotonic stimulation induced the transient-sharp ATP release pattern (Fig. 1A and Fig. 5Aa), while treatment with TGFβ (10 ng/ml, 1-3days) changed the pattern of ATP release to diffuse (Fig. 5Ab). In breast cell lines, we simultaneously cultured 3 types of cell lines on 3 separated collagen-gel patches on a cover glass and measured the ATP luminescence of each cell colony at the same time (Fig. 5Ba). The peak intensity of ATP luminescence induced by S1P (1 μM) and the subsequent application of hypotonic solution (50%) was measured. In all tested breast cell lines, TGFβ treatment enhanced the diffuse ATP releases induced by both S1P and hypotonic solution (Fig. 5Bb, summarized data from 3 types of cell lines), although the degree of enhancement somewhat differed among the cell lines (Fig. S2A).

**Fig. 5.**
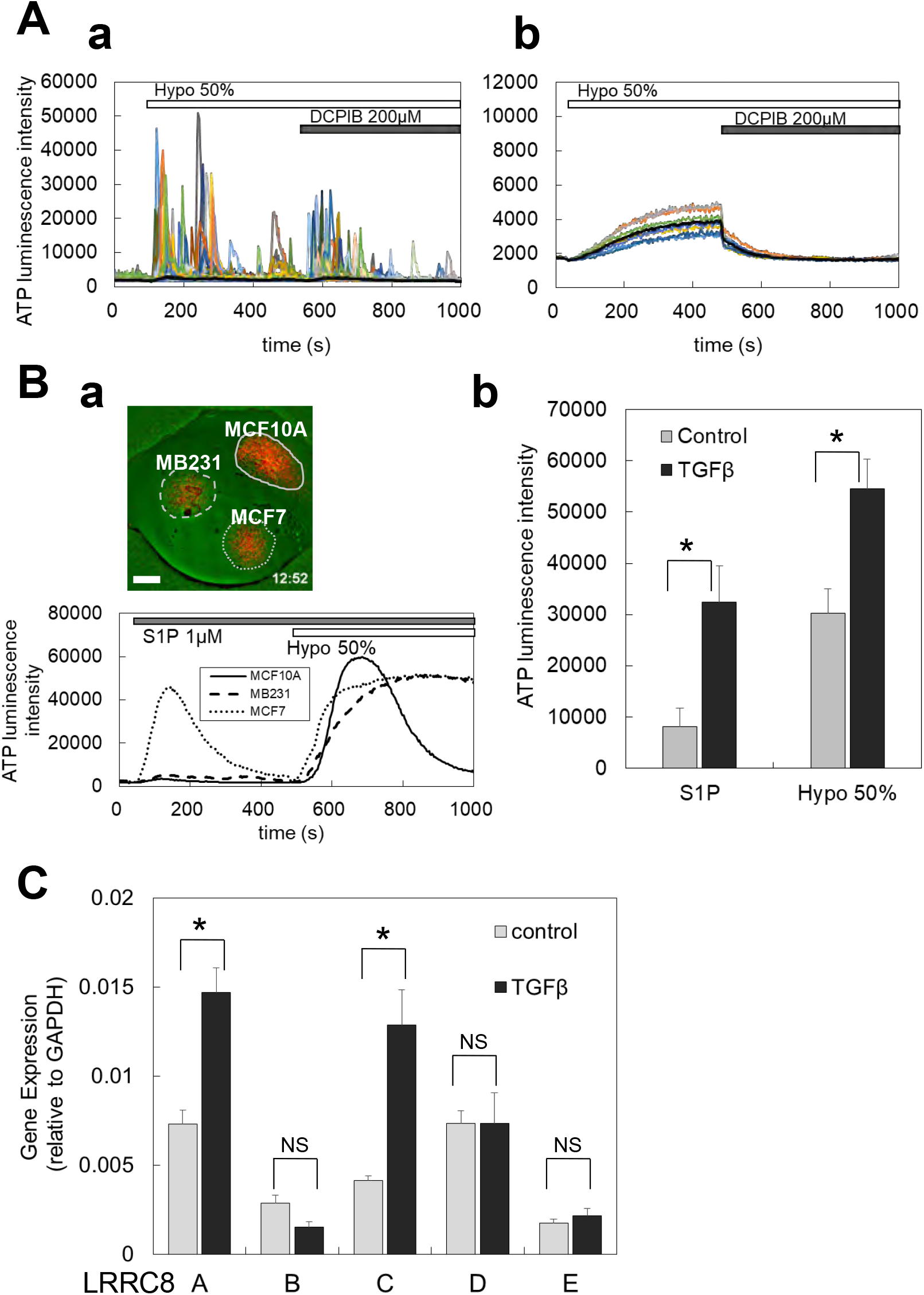
TGFβ treatment induced or enhanced the diffuse ATP release pattern and increased the expression of LRRC8 isoforms. **A:** In primary cultured mammary epithelial cells, TGFβ treatment changed the ATP release pattern from transient-sharp to diffuse. **a,** In primary cultured cells, the transient-sharp ATP release was usually induced by hypotonic stimulation (50%), and DCPIB did not block, rather it enhanced, the transient-sharp ATP release. **b,** After TGFβ treatment (10 ng/ml, 2 days), the release of ATP by hypotonic stress changed to the diffuse pattern, and DCPIB blocked the release of ATP. **B:** In breast cell lines, TGFβ enhanced the diffuse ATP release induced by both S1P and hypotonic stress. **a,** Three types of breast cell lines (MCF10A, MDA-MB231, MCF7) cultured on 3 collagen-gel patches each on a 22-mmφ cover glass placed in a small perfusion chamber. This shows an example of the measurement of the ATP release using this system. Scale bar is 2 mm. In lower panel, the time courses of the typical ATP response induced by S1P (1 μM) and subsequently applied hypotonic solution (50%) are shown. **b,** The Peak intensity of ATP luminescence induced by S1P and hypotonic solution was measured in each cell line in control or TGFβ-treated cells (shown in Fig. S2A). Plot of the average of data from 3 cell lines shows that TGFβ treatment enhanced the diffuse release of ATP by both S1P and hypotonic stress. Mean + S.E. (N=12), t-test; *: p<0.001. **C:** The LRRC8 isoforms expressed in rbeast cell lines and TGFβ treatment enhanced the expression. The expression of LRRC8 isoforms (A, B, C, D, E) in control and TGFβ-treated cells was measured with RT-qPCR and normalized to the GAPDH within each sample. The data from 3 cell lines were averaged. Differences among the cell lines are shown in Fig. S2B. Mean + S.E. (N=9), t-test; NS: no significance, *: p<0.01.

### LRRC8A isoforms were expressed in breast cell lines, and TGFβ enhanced the expression

The molecular entity of VRACs has recently been revealed as LRRC8, which shows 5 isoforms (A to E) (Voss FK et al., 2014; Qiu Z et al., 2014) forming a hexametric heteromer, although there are still points of contention, such as the contribution of each isoform to the channel functions of VRACs (Jentsch TJ et al., 2016). We checked whether the LRRC8 isoforms (A, B, C, D, and E) were expressed in breast cell lines using RT-qPCR. In control cultures, every isoform was expressed (Fig. 5C). Among them, LRRC8A, C and D were prominent, while LRRC8B and E were not very prominent. After the TGFβ treatment, the expressions levels of LRRC8A and C were further enhanced (Fig. 5C). There were some differences in the degree of the enhancement among the cell lines. The degree of enhancement somewhat differed among the cell lines (Fig. S2B).

### Gene silencing with shRNA for LRRC8A

To confirm the contribution of VRACs to the ATP release pathway of the diffuse pattern, we next knocked down LRRC8A, an indispensable subunit of VRACs, using shRNA. LRRC8A silencing with shRNA was performed using a retrovirus mediated gene transfer. Two shLRRC8A vectors (shA1 and shA2), and a non-targeting control vector (NTControl) expressed cells were cloned in three breast cell lines. RT-qPCR of the LRRC8 isoforms (A to E) showed that both shLRRC8A1 and shLRRC8A2 vectors suppressed the expression of LRRC8A but not other isoforms (B, C, D, E) in the average of three cell lines (Fig. 6A) and in each cell line: MDA-MB231, MCF7 and MCF10A (Fig. S3).

**Fig. 6.**
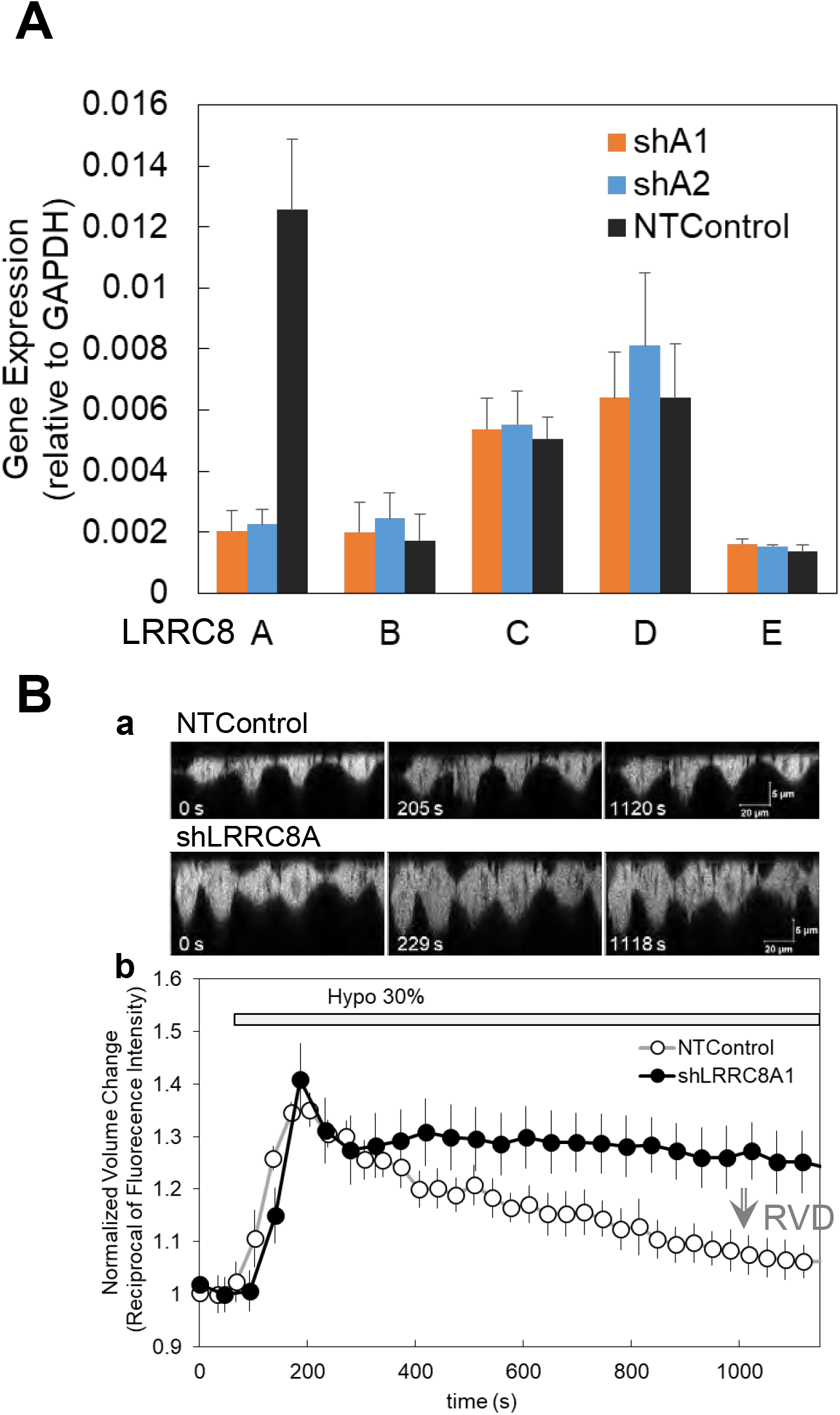
Gene silencing with shRNA for LRRC8A and suppression of the regulatory volume decrease (RVD) in knock-down cells. **A:** Two types of shRNA for LRRC8A (shA1 and shA2) suppressed the LRRC8A gene expression but did not suppress the expression of other LRRC8 isoforms (LRRC8B to E) in breast cell lines. Average of the data from 3 cell lines is shown. Differences among the cell lines are shown in Fig. S3. The expression of LRRC8 isoforms (A, B, C, D, E) in the cells treated with non-targeting control (NTControl), shA1 and shA2 were measured by RT-qPCR and normalized to the GAPDH within each sample. **B:** The knock down of LRRC8A suppressed the regulatory volume decrease (RVD) after the induction of volume increase by hypotonic stress. **a,** A time series of X-Z cross-section of MDA-MB231 cells, which expressed GFP in cytosol, was monitored with a confocal microscope. upper: NTControl, lower: shLRRC8A treated cells. **b,** The time course of the cell volume changes induced by hypotonic stress, which was calculated by the reciprocal of the fluorescence intensity and normalized to the initial value. Control cells (open circle) showed a volume increase following a volume decrease after hypotonic stress, which means RVD. RVD was suppressed in LRRC8A1 knock-down cells (closed circle).

We first checked if the deletion of LRRC8A affected an essential VRAC function in the cells—regulatory volume decrease (RVD)—after an increase of volume by hypotonic stress. Cell volume changes were estimated from changes of fluorescence intensity in a certain unit area of a perpendicular cross-section image obtained by a confocal X-Z-T scan. Hypotonic stress (30%) caused the cell height, which was observed in a perpendicular cross-section image, to increase in both shLRRC8A1 and non-targeting control cells at approximately 200 s, and decrease in control cells; however, it remained unchanged in shLRRC8A1 cells at approximately 1000s (Fig. 6Ba). The cell volume is in inverse proportion to the fluorescence intensity of the cells. The cell volume change calculated by the reciprocal of fluorescence intensity was rapidly increased until 200 s in both control and shLRRC8A1 cells (Fig. 6Bb). Thereafter, the cell volume was gradually decreased in control cells, this meant a normal RVD, whereas the cell volume was kept constant in shLRRC8A1 cells (Fig. 6Bb). This result showed that the deletion of LRRC8A suppressed the RVD function in the cells.

### LRRC8A knockdown suppressed the diffuse release of ATP induced by both hypotonic stress and S1P

The ATP-release induced by hypotonic stress was measured in the cells transfected with two sh LRRC8A vectors (shA1 and shA2) and NTControl. Three types of cells (shA1, shA2 and NTControl) were cultured on a cover glass and the ATP luminescence changes induced by 30% and 50% hypotonic solutions in each cell line were simultaneously measured. The application of hypotonic solution evoked an apparent release of ATP in NTControl cells but not in shA1 or shA2 cells (MCF10A, Fig. 6A, Suppl Movie 6). All data on the suppressive effects of LRRC8A deletion on the peak intensity of the ATP release in 3 cell lines are summarized in Fig. 7B (30% Hypo) and 7C (50% Hypo). shA1 and shA2 suppressed the release of ATP by 74% and 64% (30% Hypo), respectively, and 88% and 82% (50% Hypo), respectively.

**Fig. 7.**
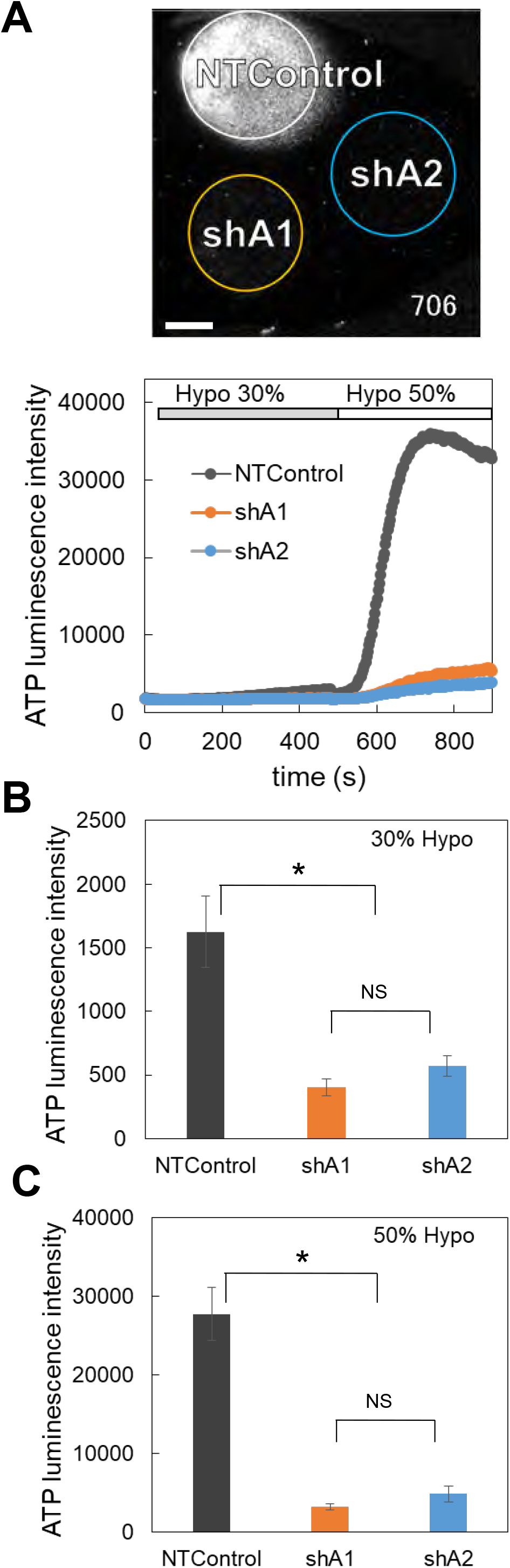
The knock-down of LRRC8A suppressed the diffuse ATP release induced by hypotonic stress. **A:** ATP release induced by hypotonic stress (30% and 50%, applied sequentially) was measured in the cells transfected with two shLRRC8A vectors (shA1 and shA2) and NTControl simultaneously, which were cultured on a cover glass (upper image and Supplemental Movie 6). Scale is 2 mm. Lower trace is the time course of measured luminescence intensity. An example of MCF10A cells. Other cell lines (MCF7, MDA-MB231) showed similar results. **B, C:** The suppression effects of shA1 and shA2 on the ATP release induced by hypotonic stress (D: 30% and E: 50%) were summarized using all data from each cell line. Mean ± S.E. (N=15), t-test; *: p<0.001.

The deletion of LRRC8A by both shRNAs also suppressed the S1P-induced release of ATP (MCF7, Fig. 8A, and Supplemental Movie 7). All data on the suppressive effects of shA1 and shA2 on the S1P-induced release of ATP in 3 cell lines are summarized in Fig. 8B. Both shA1 and shA2 suppressed the ATP release by 86%.

**Fig. 8.**
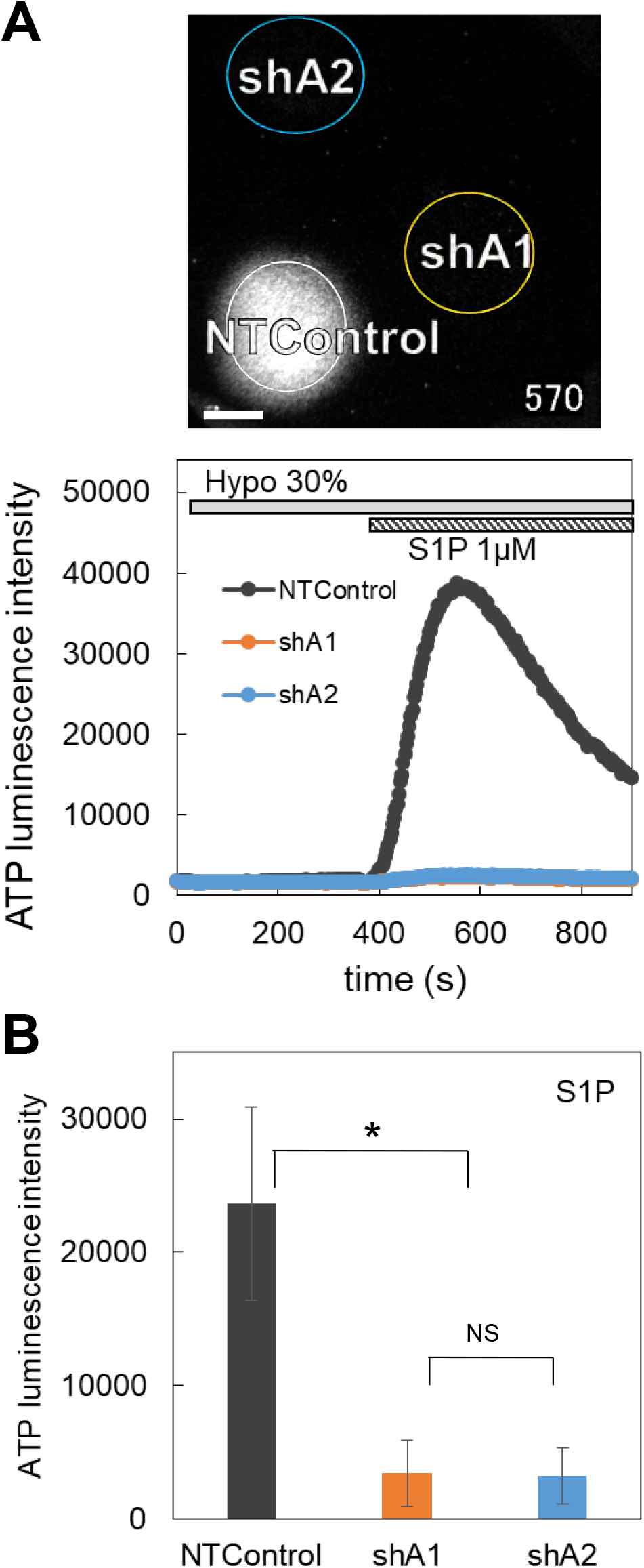
The knock down of LRRC8A suppressed the diffuse ATP release induced by the application of S1P. **A:** ATP release induced by by S1P (1 μM) subsequent to 30% hypotonic stress was measured in the cells (MCF7) transfected with two shLRRC8A vectors (shA1 and shA2) and NTControl simultaneously, which were cultured on a cover glass (upper image and Supplemental Movie 7). Scale is 2 mm. An example of MCF7 cells. Lower trace is the time course of measured luminescence intensity. Other breast cell lines (MCF10A, MDA-MB231) showed similar results. **B:** The suppression effects of shA1 and shA2 on the ATP release induced by S1P were summarized using all data from each cell line. Mean ± S.E. (N=6), t-test; *: p<0.01.

## Discussion

We herein revealed the existence of two different patterns of ATP release following hypotonic stress in mammary epithelial cells by our real time ATP imaging study and clarified the mechanism underlying the induction of diffuse-slow but substantial and prolonged pattern of ATP release in breast cell lines. The diffuse ATP release pattern was blocked by DCPIB, and suppressed by the knockdown of LRRC8A using shRNA. Thus, it was concluded that VRACs contributed to the diffuse release of ATP by hypotonic stress in breast cell lines. VRACs facilitate the passage of various substances, including ATP (Hisadome K et al., 2002; Burow P et al., 2015; Gaitán-Peñas H et al., 2016), glycine, aspartate, glutamate, GABA, taurine, myo-inositol and lactate in addition to Cl^−^ (Lutter D et al., 2017; Schober AL et al., 2017). It is debatable whether or not VRACs are a pathway of ATP release (Sabirov RZ and Okada Y, 2005; Liu HT et al., 2009). However, it is now clear that a variation of heteromeric subunit structures in the LRRC8 hexamer are related to these variations in the properties of VRACs (Planells-Cases, R et al., 2015; Gaitán-Peñas H et al., 2016; Jentsch TJ et al., 2016; Okada T et al., 2017). In addition to the indispensable component LRRC8A, the expression of LRRC8D seems to be important for the permeability of large solutes (Planells-Cases, R et al., 2015; Lutter D et al., 2017; Schober AL et al., 2017), whereas LRRC8C appears to be related to the permeability of charged osmolytes (Schober AL et al., 2017); these are supported by cryo-EM structure studies (Deneka et al., 2018; Nakamura, R et al., 2020). Recently it was reported that LRRC8C promoted cGAMP (2’3’-cyclic-GMP-AMP) transport, whereas this was inhibited by LRRC8D (Lahey LJ et al., 2020). There is still ambiguity in the role of each subunit and their combinations. Our results suggest that in addition to LRRC8A LRRC8C and D are important subunits for the release of ATP, given its markedly high expression and TGFβ-dependent increases in breast cell lines (Fig. 5C, Fig. S2B).

VRACs are not only a key element of vertebrate cell volume regulation but also participates in various physiological and pathophysiological processes (Okada Y et al., 2009; Pedersen SF et al., 2016). In human cervical cancer cells, the inhibition of VRAC resulted in G0/G1 arrest (Shen MR et al., 2000). In HeLa cells, the expression of VRAC was positively correlated with the rate of cell migration during cell cycle progression (Mao J et al., 2009). The VRAC activity in cancerous nasopharyngeal epithelial cells was much larger than that in normal human nasopharyngeal epithelial cells, consistent with the growth ability (Zhu L et al., 2012). The expression of LRRC8A was elevated in the tissues of colorectal cancer patient and shortened their survival time (Zhang H et al., 2018). Thus VRACs are implicated in the development and progression of cancer, and are a potential cancer drug target (Rong Xu et al., 2020).

VRACs were isovolumetrically activated by intracellular GTPγS, purinergic signaling, bradykinin, mGluR, ROS and Ca^2+^ signaling (Pedersen SF et al., 2016, Okada Y et al., 2019, Bertelli S et al., 2021), as well as by a decrease in ionic strength (Syeda R et al., 2016). S1P also induced the diffuse release of ATP isovolumetrically, and this was blocked by DCPIB and suppressed in LRRC8A-knockdown cells (Fig. 4, Fig. 8). This shows that S1P activated VRACs and induced the release of ATP. Interestingly, S1P induced release of ATP enhanced after hypotonic stress (Fig. 4D), implying that S1P receptors on the membrane folding, such as caveolae, might appear during the cell expansion by the hypotonic stress. S1P induced ATP release via VRAC formed an autocrine link between inflammatory sphingolipid and purinergic signaling in macrophages (Burow P et al., 2015) and microglia (Zahiri D et al., 2021). S1P is an inflammatory mediator and is produced by sphingosine kinase, which is activated by several inflammatory signaling molecules, including bacterial b b lipopolysaccharide (LPS), PDGF, TNFα, thrombin, IgE-bound antigen and ATP (Burow P et al., 2015; Burow P and Markwardt F, 2014). S1P is rich in the cancer microenvironment (Nagahashi M et al. 2012) and plays important roles in cancer progression via diverse pathways of its G-protein coupled receptors, which implicates S1P pathway as a therapeutic target (Pyne NJ and Pyne S, 2010; Ogretmen B, 2018; Nagahashi M et al., 2018). It is plausible that the induction of the release of ATP via VRAC is an important function of S1P in the cancer microenvironment (Furuya K et al., 2021).

In addition to the diffuse ATP release pattern induced via VRACs, the transient-sharp ATP release pattern was induced by hypotonic stress in mammary epithelial cells (Fig. 1A, Fig. 2B). The transient-sharp pattern usually appeared in primary cultured cells and the diffuse ATP release pattern was observed in breast cell lines, although both patterns co-existed, to different degree, in the cells. Sometimes spontaneous release of ATP with transient-sharp pattern was observed without any stimulation. In breast cell lines, cholera toxin treatment caused the diffuse ATP release pattern to disappear and induced the transient-sharp pattern (Fig. 2). The transient-sharp pattern was not suppressed by DCPIB treatment (Fig. 3C, Fig. 5Aa), nor was it suppressed in LRRC8A-knockdown cells. We tested several other blockers, CBX and 10PANX for hemi channels, NPPB for Cl^−^ channels, CFTR(inh)-172 for CFTR, and a cocktail of brefeldin A, monensin and NEM, and clodronate for exocytosis, but we noted no obvious effect on the transient-sharp pattern. As such, the pathway of the transient-sharp ATP release remains unclear at present. However, the transient-sharp ATP release is so prominent in primary cultured or differentiated cells that it may contribute certain cell functions, which remains to be elucidated.

Cholera toxin is produced by *Vibrio cholerae* and is a multifunctional protein that influences various cells, including the immune system, and which also acts as an anti-inflammatory agent (Beharati K and Ganguly NK, 2011; Baldauf K et al., 2015). Cholera toxin was shown to suppress carcinogenesis in colon cancer (Doulberis M et al., 2015) and may exert contact inhibition by binding to glycosphingolipids in MCF10A cells (Huang X et al., 2017). One of the functions of cholera toxin is to activate adenylate cyclase, which increases the intracellular cAMP level. Cholera toxin also exerts its activity through the non-toxic cholera toxin B subunit that specifically binds to the surface receptor GM1 ganglioside on lipid rafts (Baldauf K et al., 2015; Day CA and Kenworthy AK, 2015) and works as a raft cross-linker or triggers endocytosis of the binding area. Our preliminary result, that cholera toxin B subunit mimicked the effect but dibutyryl-cAMP treatment did not induce the effect suggested the latter case. Interestingly, GM1 is also a sphingosine kinase activator that induces the production of S1P (Wang F et al., 1996).

In contrast to the effect of cholera toxin, TGFβ treatment converted the transient-sharp ATP release to the diffuse pattern in primary cultured mammary epithelial cells (Fig. 5A) and enhanced the diffuse pattern in breast cell lines (Fig. 5B). TGFβ, which was first implicated in mammary epithelial development, is critically important for mammary morphogenesis and the secretory function (Moses H and Barcellos-Hoff MH, 2011; Daniel CW et al., 1989). TGFβ is also known to exist abundantly at tumor sites and plays central roles in carcinogenesis (Syed V, 2016). TGFβ signaling helps to regulate crucial cellular activities, such as cell growth, differentiation, apoptosis, motility, invasion, extracellular matrix production, angiogenesis and the immune response. However, the role and signaling pathway of TGFβ are totally cell-context dependent. TGFβ is an important inducer of the EMT in both development and carcinogenesis (Syed V, 2016; Martínez-Ramírez AS et al., 2017). There have been some reports of TGFβ inducing the EMT in mammary epithelial cell lines, including MCF10A (Zhang J et al., 2014), MDA-MB231 and MCF7 (Romagnoli M et al., 2012; Chen L et al., 2017). While the mechanism by which TGFβ influences the release of ATP via VRACs is unclear, the EMT may be associated with the change in the pattern of ATP release.

In the present study, we used three breast cell lines that originated from different tissue states. MDA-MB231 cells were derived from adenocarcinoma and are highly aggressive with triple-negative properties (Cailleau R et al., 1974). MCF7 cells were a ductal carcinoma cell line (Soule HD et al., 1973). MCF10A cells originated from benign tumors of fibrocystic disease and are non-carcinogenic, although not normal karyotypically (Soule HD et al., 1990). These cell lines possess different characteristics, as demonstrated by gene and protein expression profiling (Charafe-Jauffret E, et al., 2006). However, all of these cell lines are immortal and show undifferentiated properties in usual culture conditions. Treatment with cholera toxin, which suppresses inflammation and carcinogenesis, and which occasionally induces differentiation in various types of cells, suppressed the diffuse ATP release pattern and induced the transient-sharp release pattern. Treatment with TGFβ, which sometimes induced carcinogenesis and the EMT, induced the diffuse ATP release pattern in primary cultured cells and enhanced the diffuse ATP release pattern in breast cell lines. In addition to mammary cell lines, a lung cancer cell line (A549) showed a similar diffuse pattern (Furuya K et al., 2014). The diffuse ATP release pattern was rarely observed in primary cultured cells of mammary glands and subepithelial fibroblasts in the intestine. These results suggest that the appearance of diffuse ATP release depends on the undifferentiated state of the cells, including cancer cells. The slow and diffuse, but substantial and prolonged release of ATP via VRACs was demonstrated as a source of ATP—and accordingly adenosine—in the cancer microenvironment.

VRACs are differentially regulated through the cell cycle, and the inhibition of VRACs suppresses cell proliferation. In cancer cells, VRAC regulation is tightly coupled with migration, proliferation and metastasis. Although these mechanisms of regulation have not been elucidated, the release of ATP via VRACs in undifferentiated cells—revealed for the first time in this study—certainly plays an important role in the functions of the VRAC in cancer. The VRACs are known to be activated not only by hypotonic stress but also in physiological and pathophysiological processes, including cancer, edema, cell proliferation, migration, angiogenesis and apoptosis. These features imply the suitability of VRACs as a new therapeutic target in the cancer microenvironment.

## Grants

This work was supported by a grant for collaborative research between Nagoya University and R-Pharm (2614Dj-02b) and by JSPS KAKENHI Grants nos. 24590274, 15K08174, 18K06851 (K. F).

## Disclosures

The authors declare no conflicts of interest in association with the present study.

## Ethical approval

Animal care and all experimental procedures used in the study were approved by the Animal Experiment Review Committee, Graduate School of Medicine, Nagoya University (approval nos. 28295, 29271, 30277 and 20294).

## Supplemental Movies

**Movie 1.** The release of ATP induced by 30% hypotonic stress in primary mammary epithelial cells, shown in 3D intensity profile.

**Movie 2.** The release of ATP induced by 30% hypotonic stress in MDB-MB231 breast cancer cells, shown in 3D intensity profile.

**Movie 3.** The release of ATP induced by 30% hypotonic stress in non-treated (Control) MCF10A cells (red), overlaid on the infrared transmission image (green).

**Movie 4.** The release of ATP induced by 30% hypotonic stress in cholera toxin-treated MCF10A cells, overlaid on the infrared transmission image (green).

**Movie 5.** The release of ATP induced by 50% hypotonic stress after DCPIB treatment in cholera toxin-treated MCF10A cells.

**Movie 6.** The release of ATP induced by 50% hypotonic stress was suppressed in LRRC8A knock-down MCF10A cells (shA1 and shA2) but not in control (NTC).

**Movie 7.** The release of ATP induced by S1P (1 μM) was suppressed in LRRC8A knock-down MCF7 cells (shA1 and shA2) but not in control (NTC).

---

**Fig. S1.**
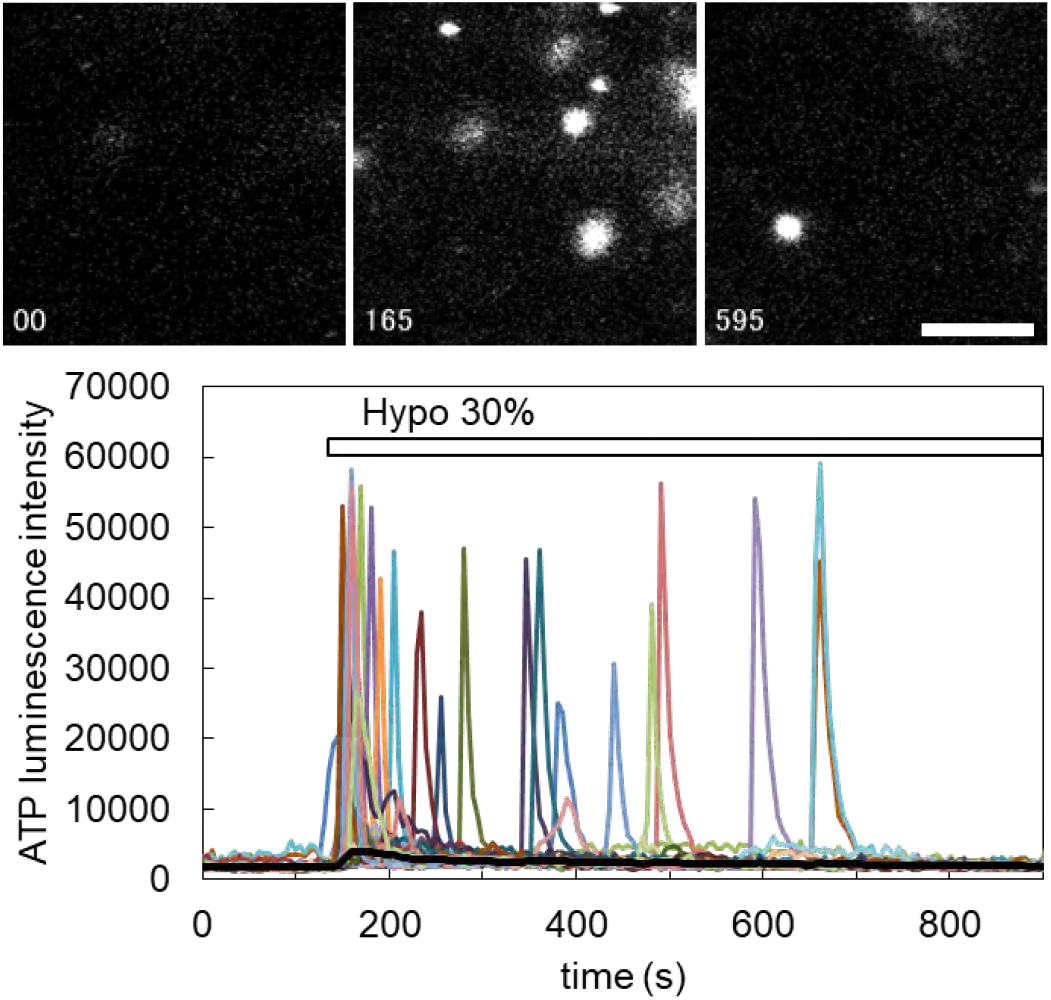
Primary culture of subepithelial fibroblasts in rat intestinal villi released ATP with transient-sharp pattern in response to 30% hypotonic stress. Subepithelial fibroblasts were isolated and cultured from rat intestinal villi by the method described in Furuya S, Furuya K (2007), and treated with endothelin 1 100 nM, which made the cells differentiate as myofibloblasts. Scale bar represents 1 mm.

**Fig. S2.**
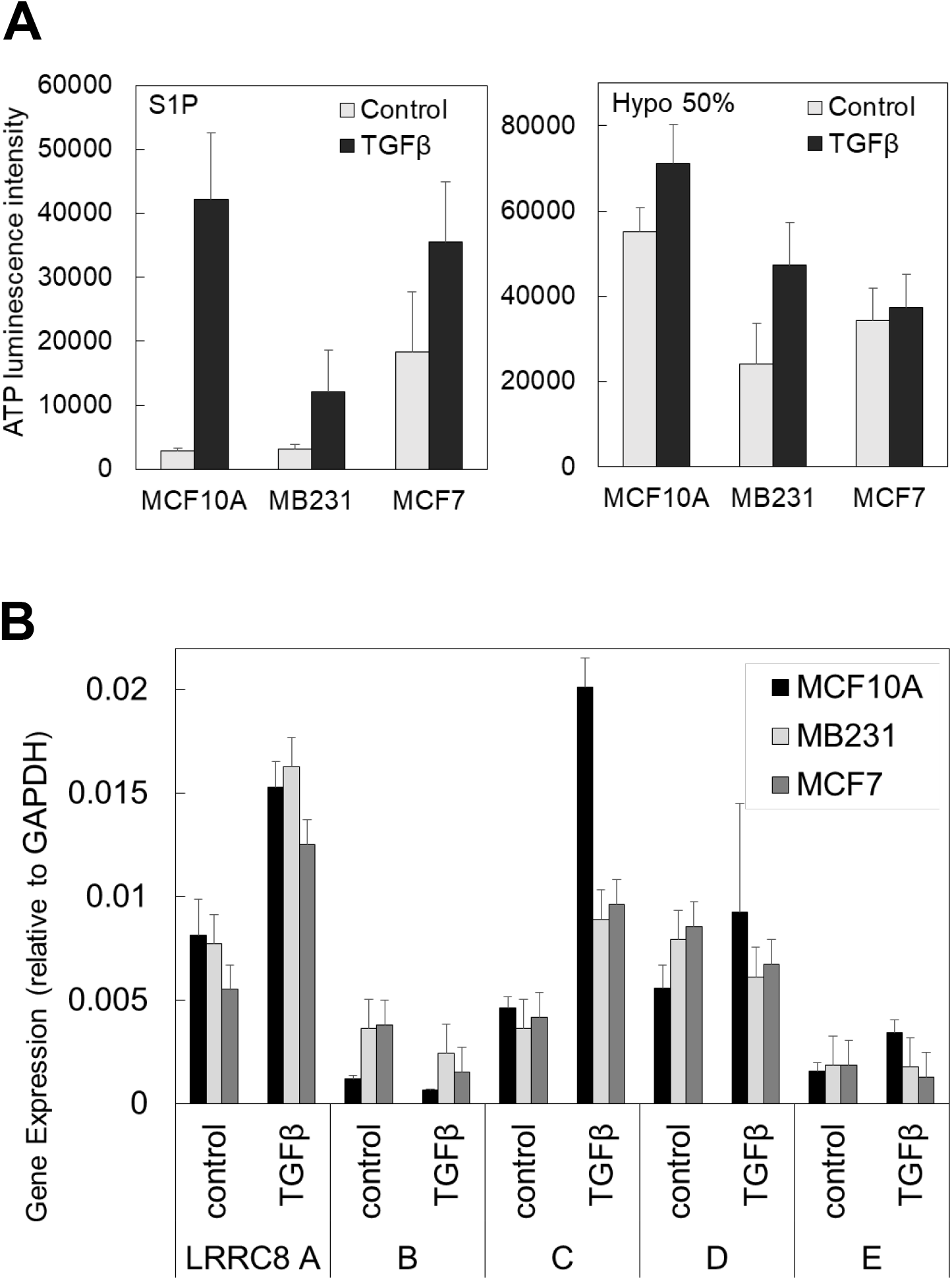
**A:** TGFβ treatment enhanced the diffuse release of ATP induced by both S1P and hypotonic stress in each breast cell line. **B:** TGFβ treatment enhanced the expression of LRRC8 isoforms in each breast cell line. Increases in LRRC8A and C were prominent.

**Fig. S3.**
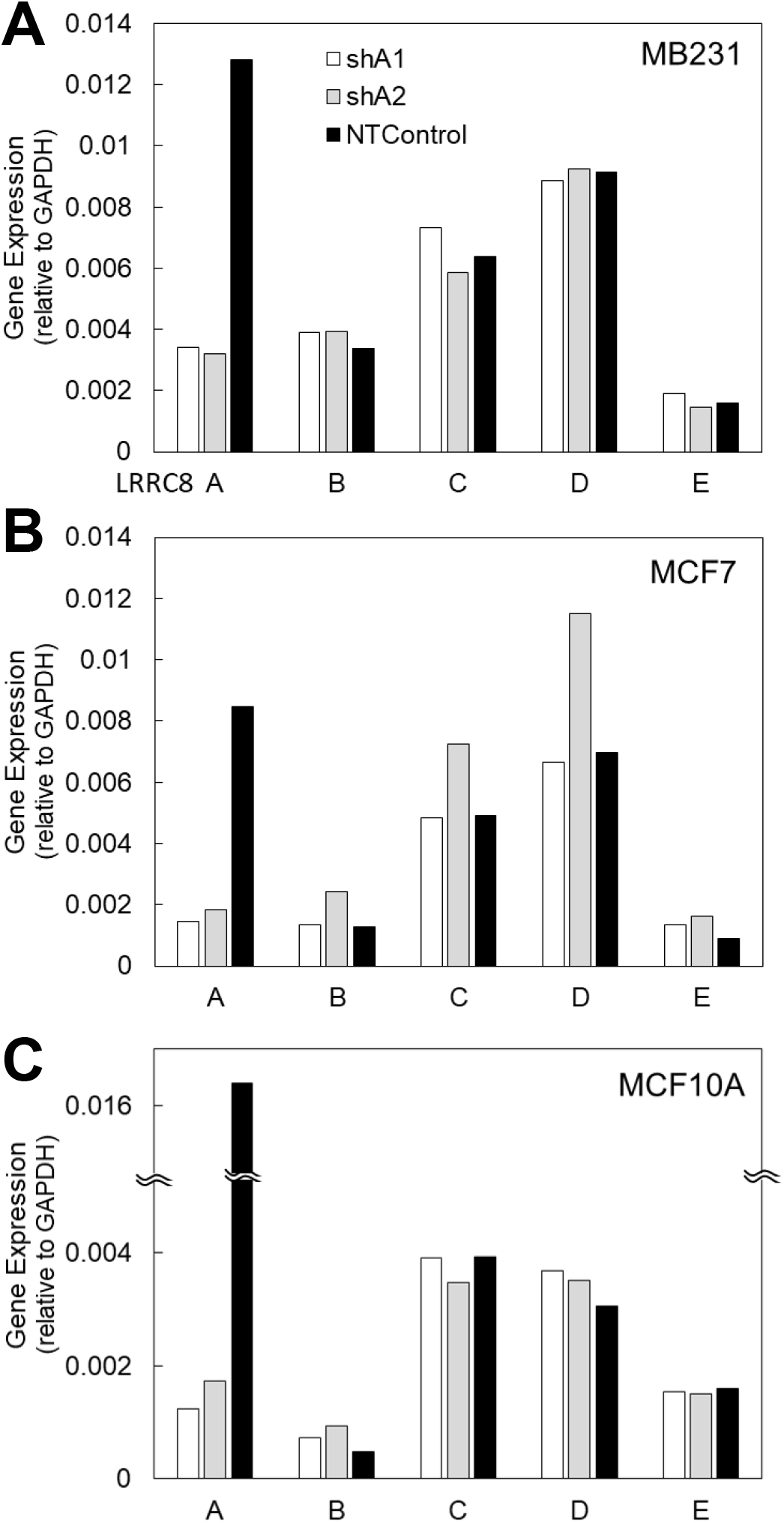
Suppression of gene expression of LRRC8A by 2 types of shRNA (shA1 and shA2) in each breast cell line. **A:** MDA-MB231, **B:** MCF7, **C:** MCF10A.

**Supplemental Table S1.**
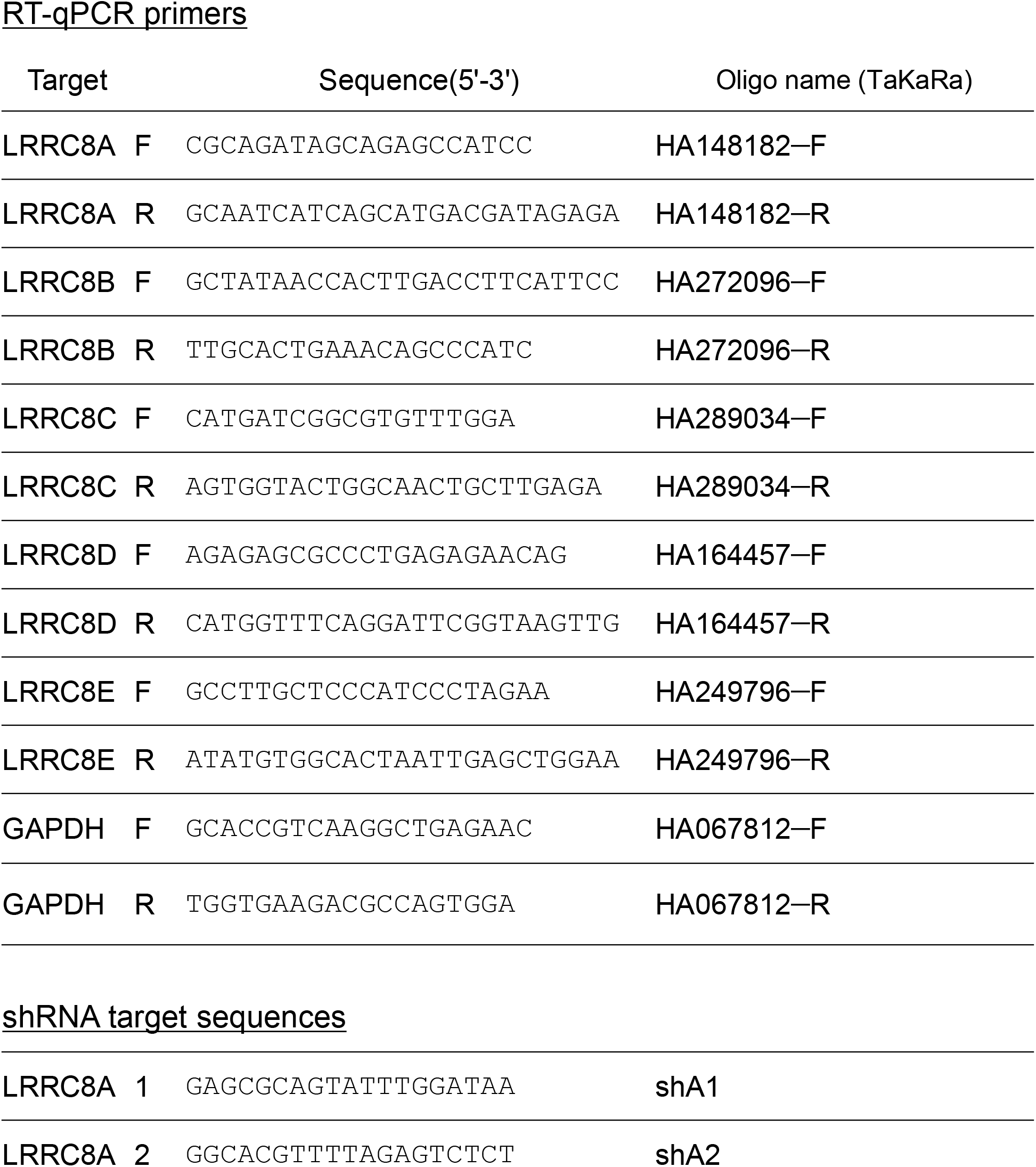
RT-qPCR primers and shRNA target sequences

## Notes

### Competing Interest Statement

The authors have declared no competing interest.

### Summary of Updates

Some sentences were added to improve an understanding of this experiment. We also corrected several mistakes in the text. No changes in Figures or data.

## References

Abascal F, Zardoya R (2012) LRRC8 proteins share a common ancestor with pannexins, and may form hexameric channels involved in cell-cell communication. Bioessays 34:551–560. doi:10.1002/bies.201100173.

Antonioli L, Blandizzi, C, Pacher P, Hasko G (2013a) Immunity, inflammation and cancer: a leading role for adenosine. Nature Rev Cancer 13: 842–857.

Antonioli L, Pacher P, Vizi ES, Hasko G (2013b) CD39 and CD73 in immunity and inflammation. Trends Mol Med 19:355–367.

Baldauf K, Royal JM, Hamorsky KT, Matoba N (2015) Cholera Toxin B: One Subunit with Many Pharmaceutical Applications. Toxins 7:974–996.

Beavis PA, Milenkovski N, Henderson MA, Darcy PK, et al. (2015a) Adenosine Receptor 2A Blockade Increases the Efficacy of Anti–PD-1 through Enhanced Antitumor T-cell Responses. Cancer Immunol Res 3: 506–517.

Beavis PA, Slaney CY, Milenkovski N, Henderson MA, Darcy PK et al. (2015b) CD73:A potential biomarker for anti-PD-1 therapy. OncoImmunology 4: 1–5.

Bharati K, Ganguly NK (2011) Cholera toxin: A paradigm of a multifunctional protein. Indian J Med Res 133: 179–187.

Bertelli S, Remigante A, Zuccolini P, Barbieri R, Ferrera L, Picco C, Gavazzo P, Pusch M (2021) Mechanisms of Activation of LRRC8 Volume Regulated Anion Channels. Cell Physiol Biochem: 55(S1):41–56. doi: 10.33594/00000032.

Burnstock G, Virgilio FDi (2013) Purinergic Signalling and Cancer. Purinergic Signalling 9:491–540. Doi: 10.1007/s11302-013-9372-5.

Burow P, Markwardt F (2014) When S1P meets ATP. Channels 8: 385–386.

Burow P, Klapperstück M, Markwardt F (2015) Activation of ATP secretion via volume-regulated anion channels by sphingosine-1-phosphate in RAW macrophages. Pflügers Archiv-Eur J Physiol 467:1215–1226.

Cailleau R, Young R, Olive M, Reeves WJ Jr (1974) Breast Tumor Cell Lines From Pleural Effusions. J Natl Cancer Inst 53: 661–674.

Charafe-Jauffret E, Ginestier C, Monville F, Finetti P, Adélaïde J, Cervera N, Fekairi S, Xerri L, Jacquemier J, Birnbaum D, Bertucci, F (2006) Gene expression profiling of breast cell lines identifies potential new basal markers. Oncogene 25: 2273–2284.

Chen L, Fu H, Luo Y, Chen L, Cheng R, Zhang N, Guo H (2017) cPLA2α mediates TGF-β-induced epithelial–mesenchymal transition in breast cancer through PI3k/Akt signaling. Cell Death and Disease 8: e2728.

Daniel CW, Silberstein GB, Van Horn K, Strickland P, Robinson S (1989) TGF-βl-induced inhibition of mouse mammary ductal growth: Developmental specificity and characterization. Dev Biol 135:20–30.

Day CA, Kenworthy AK (2015) Function of cholera toxin B-subunit as a raft cross-linker. Essays Biochem 57: 135–145.

Decher N, Lang HJ, Nilius B, Bruggemann A, Busch AE, Steinmeyer K (2001) DCPIB is a novel selective blocker of Icl, swell and prevents swelling-induced shortening of guinea-pig artrial action potential duration. Brit J Pharm 134:1467–1479.

Deneka D, Sawicka M, Lam AKM, Paulino C, Dutzler R (2018) Structure of a volume-regulated anion channel of the LRRC8 family. Nature 558: 254–259. doi:10.1038/s41586-018-0134-y

Doulberis M, Angelopoulou K, Kaldrymidou E, Tsingotjidou A, Abas Z, Erdman SE, Poutahidis T (2015) Cholera-toxin suppresses carcinogenesis in a mouse model of inflammation-driven sporadic colon cancer. Carcinogenesis 36: 280–290.

Fazzari J, Singh G (2016) Cancer-Induced Edema/Lymphedema. In Oncodynamics: Effects of Cancer Cells on the Body. Springer pp85–103.

Friard J, Tauc M, Cougnon M, Compan V, Duranton C, Rubera I (2017) Comparative Effects of Chloride Channel Inhibitors on LRRC8/VRAC-Mediated Chloride Conductance. Front Pharmacol 8: Article 328.

Furuya K, Sokabe M, Grygorczyk R (2014) Real-time Luminescence Imaging of Cellular ATP Release. Methods 66: 330–344. doi: 10.1016/j.ymeth.2013.08.007.

Furuya K, Hirata H, Kobayashi T, Sokabe M (2021) Sphingosine-1-Phosphate Induces ATP Release via Volume-Regulated Anion Channels in Breast Cell Lines. Life 11: 851. https://doi.org/10.3390/life11080851.

Furuya S, Furuya K (2007) Subepithelial fibroblasts in intestinal villi: Roles in intercellular communication. International Review of Cytology 264: 165–223. doi: 10.1016/S0074-7696(07)64004-2.

Gaitán-Peñas H, Gradogna A, Laparra-Cuervo L, Solsona C, Fernández-Dueñas V, Barrallo-Gimeno A, Ciruela F, Lakadamyali M, Pusch M, Estévez R. (2016) Investigation of LRRC8-Mediated Volume-Regulated Anion Currents in Xenopus Oocytes. Biophys J 111: 1429–1443. doi: 10.1016/j.bpj.2016.08.030.

Ginzberg MB, Kafri R, Kirschner M (2015) On being the right (cell) size. Science 48:1245075.

Hausler SF, Del Barrio IM, Diessner J, Stein RG, et al. (2014) Anti-CD39 and anti-CD73 antibodies A1 and 7G2 improve targeted therapy in ovarian cancer by blocking adenosine-dependent immune evasion. Am J Transl Res 26:129–39.

Hirata H, Samsonov M and Sokabe M (2017) Actomyosin contractility provokes contact inhibition in E-cadherin-ligated keratinocytes. Sci Rep 7: 46326. doi:10.1038/srep46326.

Hisadome K, Koyama T, Kimura C, Droogmans G, Ito Y, Oike M (2002) Volume-regulated Anion Channels Serve as an Auto/Paracrine Nucleotide Release Pathway in Aortic Endothelial Cells. J Gen Physiol 119: 511–520.

Huang X, Schurman N, Handa K, Hakomori S (2017) Functional role of glycosphingolipids in contact inhibition of growth in a human mammary epithelial cell line. FEBS Letters 591:1918–1928.

Jentsch TJ, Lutter D, Planells-Cases R, Ullrich F, Voss FK (2016) VRAC: molecular identification as LRRC8 heteromers with differential functions. Pflügers Arch - Eur J Physiol 468:385–393.

Lahey LJ, Mardjuki RE, Wen X, Hess GT, Ritchie C, Carozza JA, Böhnert V, Maduke M, Bassik MC, Li LL (2020) LRRC8A:C/E Heteromeric Channels Are Ubiquitous Transporters of cGAMP. Molecular Cell 80:578–591.e5. doi:10.1016/j.molcel.2020.10.021.

Liu HT, Akita T, Shimizu T, Sabirov RZ, Okada Y (2009) Bradykinin-induced astrocyte-neuron signalling: glutamate release is mediated by ROS-activated volume-sensitive outwardly rectifying anion channels. J Physiol 587: 2197–2209.

Lutter D, Ullrich F, Lueck JC, Kempa S, Jentsch TJ (2017) Selective transport of neurotransmitters and modulators by distinct volume-regulated LRRC8 anion channels. J Cell Sci 130: 1122–1133.

Mao J, Chen L, Xu B, Wanga L, Wangc L, et al. (2009) Volume-activated chloride channels contribute to cell-cycle-dependent regulation of HeLa cell migration. Biochem Pharmacol 77: 159–168.

Martínez-Ramírez AS, Díaz-Muñoz M, Butanda-Ochoa A, Vázquez-Cuevas FG (2017) Nucleotides and nucleoside signaling in the regulation of the epithelium to mesenchymal transition (EMT). Purinergic Signalling 13:1–12.

Mongin AA (2016) Volume-regulated anion channel—a frenemy within the brain. Pflügers Arch - Eur J Physiol 468:421–441.

Moses H, Barcellos-Hoff MH (2011) TGF-β Biology in Mammary Development and Breast Cancer. Cold Spring Harb Perspect Biol 2011;3:a003277.

Nagahashi M, Ramachandran S, Kim EY, Allegood JC, Rashid OM, Yamada A, Zhao P, Milstien S, Zhou H, Spiegel S and Takabe K (2012) Sphingosine-1-Phosphate Produced by Sphingosine Kinase 1 Promotes Breast Cancer Progression by Stimulating Angiogenesis and Lymphangiogenesis. Cancer Res 72:726–735. doi:10.1158/0008-5472.CAN-11-2167.

Nagahashi M, Abe M, Sakimura K, Takabe K and Wakai T. (2018) The role of sphingosine-1-phosphate in inflammation and cancer progression. Cancer Science 109: 3671–3678. doi:10.1111/cas.13802.

Nakamura R, Numata T, Kasuya G, Yokoyama T, Nishizawa T, Kusakizako T, Kato T, Hagino T, Dohmae N, Inoue M et al. (2020) Cryo-EM structure of the volume-regulated anion channel LRRC8D isoform identifies features important for substrate permeation. Commun Biol 3: 240. doi:10.1038/s42003-020-0951-z.

Nakano H, Furuya K, Furuya S, Yamagishi S (1997) Involvement of P2-purinergic receptors in intracellular Ca2+ responses and the contraction of mammary myoepithelial cells. Pflügers Arch-Eur J Physiol 435:1–8.

Ogretmen B (2018) Sphingolipid metabolism in cancer signalling and therapy. Nature Reviews Cancer 18: 33–50. doi:10.1038/nrc.2017.96.

Okada T, Islam Md R, Tsiferova NA, Okada Y, Sabirov RZ (2017) Specific and essential but not sufficient roles of LRRC8A in the activity of volume-sensitive outwardly rectifying anion channel (VSOR). Channels 11: 109–120.

Okada Y, Sato K, Numata T (2009) Pathophysiology and puzzles of the volume-sensitive outwardly rectifying anion channel. J Physiol 587, 2141–2149.

Okada Y, Okada T, Sato-Numata K, Islam Md R, Ando-Akatsuka Y, Numata T, Kubo M, Shimizu T, Kurbannazarova RS, Marunaka Y and Sabirov RZ (2019) Cell Volume-Activated and Volume-Correlated Anion Channels in Mammalian Cells: Their Biophysical, Molecular, and Pharmacological Properties. Pharmacological Reviews 71:49–88. doi:10.1124/pr.118.015917.

Pedersen SF, Okada Y, Nilius B (2016) Biophysics and Physiology of the Volume-Regulated Anion Channel (VRAC)/Volume-Sensitive Outwardly Rectifying Anion Channel (VSOR). Pflugers Arch - Eur J Physiol 468:371–383

Pellegatti P, Raffaghello L, Bianchi G, Piccardi F, Pistoia V, Virgilio FDi (2008) Increased Level of Extracellular ATP at Tumor Sites: In Vivo Imaging with Plasma Membrane Luciferase PLoS ONE 3: e2599. doi:10.1371/journal.pone.0002599.

Planells-Cases R, Lutter D, Guyader C, Gerhards NM, Ullrich F, Elger DA, Kucukosmanoglu A, Xu G, Voss FK, Reincke SM et al. (2015) Subunit composition of VRAC channels determines substrate specificity and cellular resistance to Pt-based anti-cancer drugs. EMBO J 34, 2993–3008. doi:10.15252/embj.201592409.

Pyne NJ and Pyne S. (2010) Sphingosine 1-phosphate and cancer. Nature Reviews Cancer 10: 489–503. doi:10.1038/nrc2875.

Qiu Z, Dubin AE, Mathur J, Tu B, Reddy K, Miraglia LJ, Reinhardt J, Orth AP, Patapoutian A (2014) SWELL1, a plasma membrane protein, is an essential component of volume-regulated anion channel. Cell 157:447–458.

Romagnoli M, Belguise K, Yu Z, Wang X, Landesman-Bollag E, Seldin DC,Chalbos D, Barillé-Nion S, Jézéquel P, Seldin ML,Sonenshein GE (2012) Epithelial-to-Mesenchymal Transition Induced by TGF-β1 is Mediated by Blimp-1–Dependent Repression of BMP-5. Cancer Res. 72:1–11.

Sabirov RZ, Okada Y (2005) ATP release via anion channels. Purinergic Signalling 1:311–328.

Schober AL, Wilson CS, Mong AA (2017) Molecular composition and heterogeneity of the LRRC8-containing swelling-activated osmolyte channels in primary rat astrocytes. J Physiol 595.22: 6939–6951. doi:10.1113/JP275053.

Shen MR, Droogmans G, Eggermont J, Voets T, Ellory JC, Nilius B (2000) Differential expression of volume-regulated anion channels during cell cycle progression of human cervical cancer cells. J Physiol 529(Pt 2):385–394.

Soule HD, Vazquez J, Long A, Albert S, Brennan M (1973) A Human Cell Line From a Pleural Effusion Derived From a Breast Carcinoma. J Natl Cancer Inst 51:1409–1416.

Soule HD, Maloney TM, Wolman SR, Peterson WD, Brenz R, McGrath CM, Russo J, Pauley RJ, Jones RF, Brooks SC (1990) Isolation and Characterization of a Spontaneously Immortalized Human Breast Epithelial Cell Line, MCF-10. Cancer Res 50:6075–6086.

Syed V (2016) TGF-β Signaling in Cancer. J Cell Biochem 117:1279–1287.

Syeda R, Qiu Z, Dubin AE, Murthy SE, Florendo MN, Patapoutian A et al. (2016) LRRC8 proteins form volume-regulated anion channels that sense ionic strength. Cell 164:499–511. doi:10.1016/j.cell.2015.12.031.

Virgilio FDi (2012) Purines, Purinergic Receptors\, and Cancer. Cancer Res 72: 5441–5447. doi:10.1158/0008-5472.CAN-12-1600.

Voss FK, Ullrich F, Munch J, Lazarow K, Lutter D, Mah N, Andrade-Navarro MA, von Kries JP, Stauber T, Jentsch TJ (2014) Identification of LRRC8 heteromers as an essential component of the volume-regulated anion channel VRAC. Science 344:634–638.

Wang F, Buckley NE, Olivera A, Goodemote KA, Su Y, Spiegel S (1996) Involvement of sphingolipids metabolites in cellular proliferation modulated by ganglioside GM1. Glycoconjugate J 13:937–945.

Xu R, Wang X, Shi C (2020) Volume-regulated anion channel as a novel cancer therapeutic target. Int J Biol Macromol 159, 570–576. doi:10.1016/j.ijbiomac.2020.05.137.

Zahiri D, Burow P, Großmann C, Müller CE, Klapperstück, Markwardt F (2021) Sphingosine-1-phosphate induces migration of microglial cells via activation of volume-sensitive anion channels, ATP secretion and activation of purinergic receptors. Biochim Biophys Acta (BBA) - Mol Cell Res 1868: 118915. doi:10.1016/j.bbamcr.2020.118915.

Zhang H, Deng Z, Zhang D, Li H, Zhang L, Niu J, Zuo W, Fu R, Fan L, Ye JH et al. (2018) High expression of leucine-rich repeat-containing 8A is indicative of a worse outcome of colon cancer patients by enhancing cancer cell growth and metastasis. Oncol Rep 40: 1275–1286. doi:10.3892/or.2018.6556.

Zhang J, Tian X-J, Zhang H, Teng Y, Li R, Bai F, Elankumaran S, Xing J (2014) TGF-β-induced epithelial-to-mesenchymal transition proceeds through stepwise activation of multiple feedback loops. Science Signaling 345:ra91.

Zhu L, Yang H, Zuo W, Yang L, Wanga L, et al. (2012) Differential expression and roles of volume-activated chloride channels in control of growth of normal and cancerous nasopharyngeal epithelial cells. Biochem Pharmacol 83: 324–334.

